# Mass spectrometry-based proteomic exploration of diverse murine macrophage cellular models

**DOI:** 10.1101/2024.04.09.588684

**Authors:** Jack Gudgeon, José Luis Marin Rubio, Frances Sidgwick, Matthias Trost

## Abstract

Immortalised cell lines analogous to their primary cell counterparts are fundamental to research, particularly when large cell numbers are required. Here we report that immortalisation of bone marrow-derived macrophages using the J2-virus resulted in the loss of a protein of interest, MSR1, in wild-type cells by an unknown mechanism. This led us to perform an in-depth mass spectrometry-based proteomic characterisation of common murine macrophage cell lines (J774A.1, RAW264.7, and BMA3.1A7), with comparison to the immortalised bone marrow-derived macrophages (iBMDMs), as well as primary BMDMs. This revealed striking differences in protein profiles associated with macrophage polarisation, phagocytosis, pathogen recognition, and IFN signalling. J774A.1 cells were determined to be the most similar to the gold standard primary BMDM model, with BMA3.1A7 cells the least similar due to the reduction in abundance of several proteins related closely to macrophage function. This comprehensive proteomic data offers valuable insights into the selection of specific macrophage cell lines for cell signalling and inflammation research.

## Introduction

Macrophages were first described by Élie Metchnikoff who primarily noted their phagocytic (Greek: phagein - to eat, kytos – cell) ability (Gordon, 2016). As antigen presenting cells (APCs) belonging to the innate immune system, they act as part of the body’s first line of defence against invading pathogens or damaging self-molecules by recognition of pathogen associated molecular patterns (PAMPs) or damage associated molecular patterns (DAMPs), respectively. Macrophages are highly plastic in nature, existing as a range of tissue-resident cells and adopting various polarisation states in response to different activating stimuli. These activation or polarisation states hold distinct gene signatures and drive varying macrophage functions. In reality, the macrophage polarisation state exists on a complex spectrum, however a more reductionist approach is often used to delineate distinct phenotypes (Murray et al., 2014). This binary nomenclature consists of M1 (classically activated or pro-inflammatory phenotype) and M2 (alternatively activated or anti-inflammatory phenotype) macrophages. M1 activation is driven primarily by stimuli such as interferons (IFN) or bacterial components such as lipopolysaccharide (LPS). These macrophages are primed for the promotion of tissue inflammation and pathogen killing by phagocytosis. While M2 activation is stimulated by cytokines such as IL-4, IL-10, and IL-13. M2 macrophages are more closely linked to immunoregulation, wound healing, and tissue remodelling, yet still have high phagocytic capacity (Murray, 2017).

Various degrees of macrophage activation, coupled with different phenotypes of macrophages throughout the body, introduces complex heterogeneity and difficulty in comparing macrophage subsets. To simplify biochemical investigations of macrophages, common cell lines or mouse strains are often used. Common murine macrophage models are used in effort to standardise investigations into innate immunity and inflammation. Primary cell models such as bone marrow-derived macrophages (BMDMs) from C57BL/6 mice are the gold standard as these cells are not transformed to allow indefinite culture and therefore better represent the *in vivo* macrophage phenotype (Assouvie et al., 2018).

However, immortal cell lines such as J774A.1, RAW264.7, and BMA3.1A7 are frequently used in cell signalling research, providing a cost-effective and ethically responsible alternative to *in vivo* or primary cell models. The use of cell lines in place of primary cells helps to address the ‘three Rs’, principles originally outlined by Russel & Birch in 1959, which advocate the replacement, reduction, and refinement of animal models in research (Russell and Burch, 1959). Both J774A.1 and RAW264.7 cells were obtained from BALB/c mice (Ralph et al., 1976; Raschke et al., 1978), whereas BMA3.1A7 cells were obtained from C57BL/6 mice (Magdalena et al., 1994). However, comparisons may be hindered by the use of different strains, as C57BL/6J macrophages exhibit a greater propensity for M1 polarisation, whereas BALB/c mice tend to acquire a more M2 phenotype when challenged with LPS (Mills et al., 2000). Consequently, employing differing strains to investigate macrophage responses to infection may result in conflicting evidence (Dill et al., 2015; Guo et al., 2015).

Furthermore, while cell lines offer an unlimited supply of biological material and are easier to use and manipulate, there is a caveat to their use: they must retain functional features close to those of their primary counterparts. As immortal cell lines are obtained through virus-mediated transformation or derived from tumours, they are susceptible to the accumulation of additional genetic alterations over successive passages. Consequently, these alterations can result in changes in phenotype and the loss of essential functions that are specific to macrophages (Kaur and Dufour, 2012). To ensure that experimental results can be reliably translated into real life physiology or pathophysiology, it is crucial to characterise and validate cell lines against primary cells. One common method used to immortalise BMDMs derived from mice employs the J2 recombinant gamma-retrovirus. The J2 virus originates from the replication-defective 3611-Moloney murine sarcoma virus (Mo-MSV) and harbours viral *raf* (*v-raf*) and viral *myc* (*v-myc*) oncogenes. Introduction of these oncogenes into BMDMs induces indefinite proliferation without the need for growth factors (Blasi et al., 1985; De Nardo et al., 2018). Cell lines generated using the J2 virus have been characterized using multiple different approaches, confirming the presence of common macrophage markers (Blasi et al., 1985), antigen presentation capability (Magdalena et al., 1994), and ability to polarize towards M1 and M2 phenotypes (Achita, 2015; Spera et al., 2021). However, comprehensive proteomics studies have not been performed.

Overall, macrophage biology is incredibly complex as phenotype can be affected by tissue residence, polarisation state, and genetic background. It is therefore imperative that these variables are considered when designing investigations and interpreting experimental results. Furthermore, following generation of a new model cell line such as the immortalised

BMDMs (iBMDMs) generated here by J2 virus infection, it is vital that characterisation steps are carried out to assess their validity as a cellular model. To this end, we report the characterisation of the commonly used murine macrophage cell lines (RAW264.7, J774A.1, and BMA3.1A7), alongside WT iBMDMs in comparison to WT C57BL/6 BMDMs. To facilitate this, comprehensive liquid chromatography tandem mass spectrometry (LC-MS/MS) data independent acquisition (DIA) was used to interrogate the proteome of these cell lines.

## Results

### Expression of Msr1 is lost after immortalisation of BMDMs

This project intended to continue studies (Govaere et al., 2022; Guo et al., 2019) into the role of Msr1, the macrophage scavenger receptor 1. MSR1 has been shown to have activity in macrophage polarisation, proinflammatory signalling, pathogen clearance, Alzheimer’s disease, atherosclerosis, non-alcoholic fatty liver disease, and cancer (Gudgeon et al., 2022). In order to reduce the number of animals needed for biochemical experiments, the project started by generating a cell line harbouring a genetic knock-out. Since standard CRISPR-CAS9 methods proved difficult due to *msr1’s* short exons, immortalised cell lines from wild-type and Msr1 knock-out (KO) BMDMs were generated using the J2 recombinant gamma-retrovirus (De Nardo et al., 2018).

WT and Msr1 KO iBMDMs were generated from C57BL/6N mice as described in materials and methods (**Figure S1**). Briefly, BMDMs were transduced with J2 virus sourced from the AMJ2-C11 cell line. The concentration of L929 conditioned medium in the bone marrow growth medium was then gradually reduced to 0% over 2-3 months. iBMDMs were confirmed to be macrophages by analysis of the cell surface receptors F4/80, CD11b, and CD11c. WT iBMDMs were confirmed to be F4/80^high^CD11b^high^CD11c^low^ (**Figure S2**). Expression of CD11c, a marker for dendritic cells, was reduced on iBMDM cells compared to WT BMDMs. This was potentially a result of the cells no longer being cultured in L929 conditioned media, as previous studies have shown that the L929 conditioned media used to culture BMDMs can induce CD11c expression (Heap et al., 2021; Rice et al., 2020).

Proteomics and flow cytometry experiments showed that Msr1 was not detectable in both WT and KO iBMDMs while it was detectable in BMDMs. We also added other macrophage/monocyte cell lines such as J774A.1 and RAW264.7 cells, which have high Msr1 expression as well as BMA3.1A7 cells which shows lower cell surface abundance (**Figure 1a/b**). To determine whether this loss of expression was driven by protein degradation or by changes at the genetic or epigenetic level, RT-qPCR analysis was performed to investigate changes in *Msr1* mRNA levels between WT BMDMs and WT iBMDMs (**Figure 1c**). This confirmed the loss of *Msr1* gene expression seen after immortalisation, indicating that a possible epigenetic alteration, such as CpG methylation, or genetic alteration, such as gammaretroviral integration into the genome, was responsible for the loss of MSR1.

**Figure 1.**
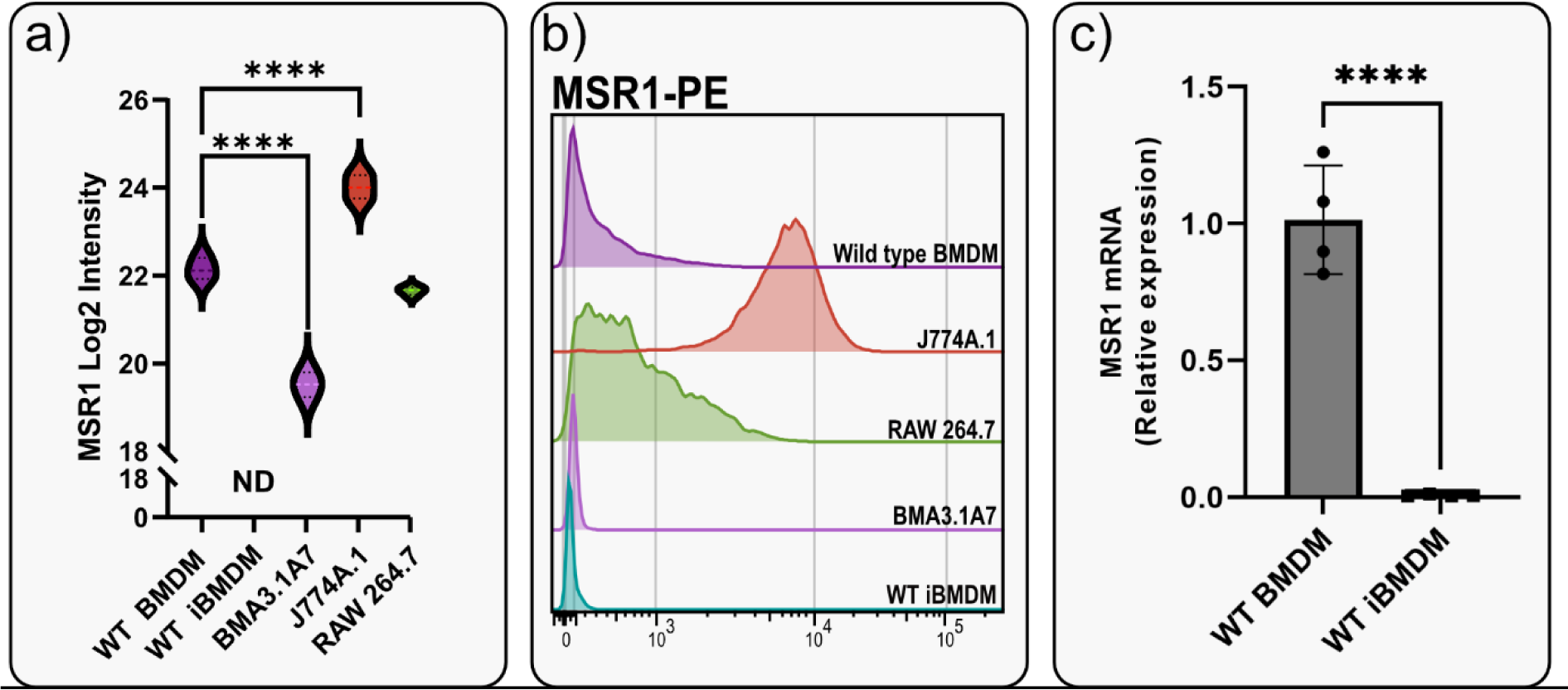
Both gene and protein expression of MSR1 was lost after J2-mediated immortalisation of BMDMs. **a)** Proteomics Log_2_ intensity data of MSR1 across all cell lines and primary macrophages (N=4). ND, not detected; ****, P-value < 0.0001 by ordinary one-way ANOVA. **b)** Flow cytometry characterisation of macrophage cell lines. The surface expression level of MSR1-PE was compared between WT BMDMs, J774A.1, RAW264.6, BMA3.1A7, and WT iBMDMs. N=1. **c)** Quantification of mRNA levels of *Msr1* in WT BMDMs and WT iBMDMs (n=4/group). Data are presented as mean ± SD (unpaired Student’s t test); ∗∗∗∗, p <0.0001.

### Proteomic characterisation of C57BL/6 iBMDMs and macrophage cell lines

Next, due to the unexpected loss of Msr1 expression in iBMDMs, we aimed to assess the suitability of cell lines for studying innate immune responses compared to primary cells, employing a proteomics approach. Using a highly sensitive liquid chromatography tandem mass spectrometry (LC-MS/MS) data-independent acquisition (DIA) method, we performed a detailed comparison between immortal macrophage cell lines (J774A.1, RAW264.6, BMA3.1A7, and iBMDMs) and primary WT BMDMs.

We identified 7971 proteins using 2h gradients on a QExactive HF mass spectrometer in DIA mode, with 6737 of the proteins being reliably identified through the detection of two or more peptides (**Table S1**). DIA-NN software was used to search against a sequence database in library-free mode. Quality control checks showed uniform dispersion and intensity distribution in all four biological replicates and groups (**Figures S3 and S4**). Moreover, principal component analysis (PCA) analysis showed that the samples were separated from each other based on their protein abundances (**Figure S5**). This suggests that there are significant differences in the protein profiles of the cells in these different groups.

Although 16 peptides for Msr1 were detected overall in cell lines, Msr1 was not detected by mass spectrometry in WT iBMDMs, validating our flow cytometry data. Furthermore, J2-immortalised BMA3.1A7 cells displayed significantly lower MSR1 protein levels compared to WT BMDMs, whereas J774A.1 cells were shown to have significantly higher MSR1 levels than WT BMDMs. RAW264.7 cells showed similar protein levels compared to BMDMs, considering that BMDMs had lower Msr1 surface levels, this might imply that in BMDMs there is a higher intracellular pool of Msr1 compared to RAW264.7 (**Figure 1a**).

Overall, as anticipated, hierarchical clustering analysis revealed distinct clustering patterns between cell lines, with WT BMDMs forming a separate cluster from cell lines (**Figure 2**). Within this distinct cluster, J774A.1 cells exhibited the closest similarity to WT BMDMs, whereas BMA3.1A7 cells displayed the least similarity in terms of proteome. RAW267.4 and iBMDMs cluster most closely together, likely because both were immortalised through the use of viruses. Furthermore, heat map analysis was employed to depict relative protein abundance, which was divided into twelve distinct clusters based on hierarchical clustering of significantly different proteins.

**Figure 2.**
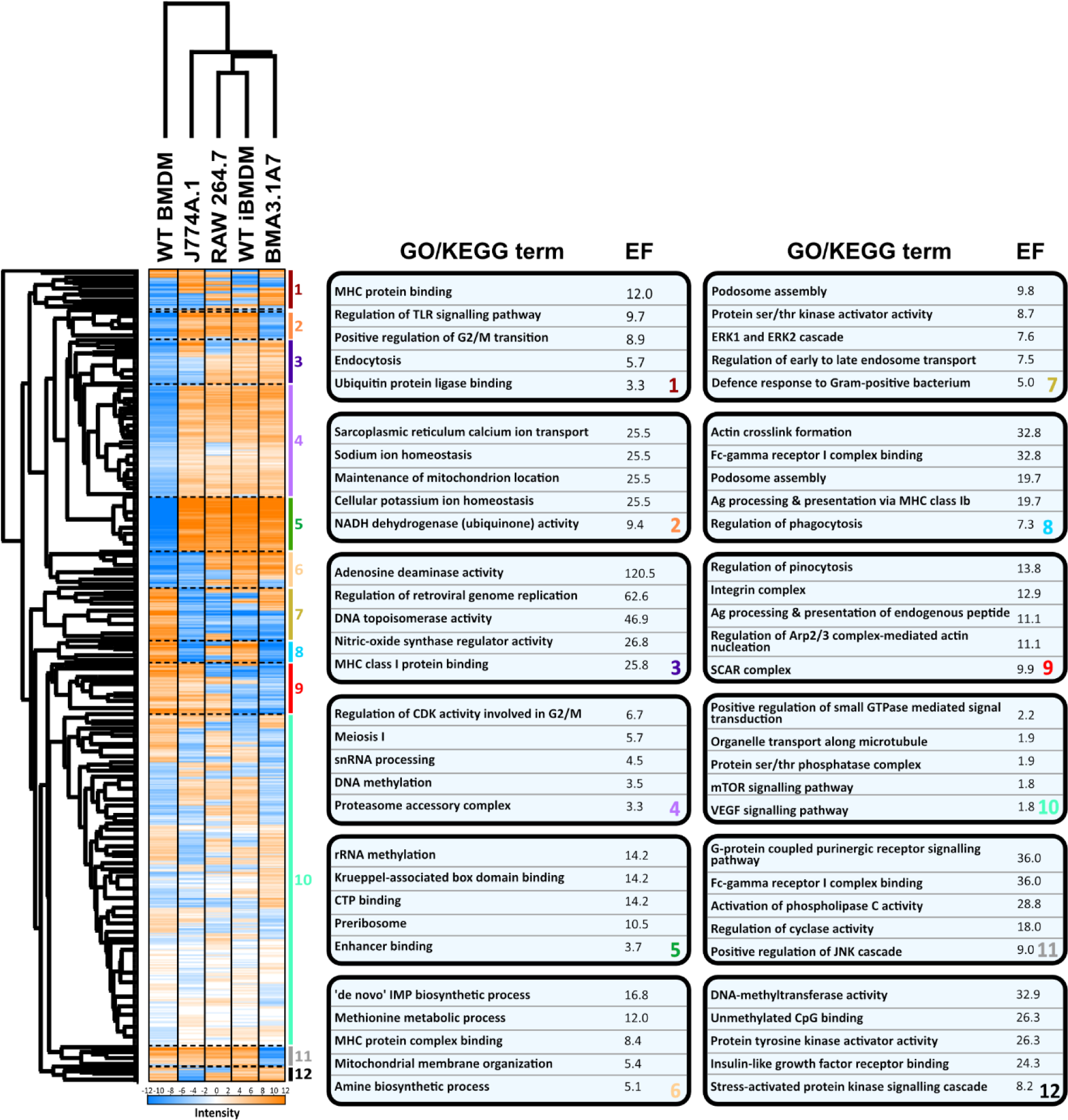
Hierarchical heatmap analysis shows the variation between primary macrophage cells and cell lines, highlighting the enriched GO or KEGG terms in each specific cluster. Proteins were normalised by Z-score, filtered for at least 3 valid values in each group, processed using ANOVA statistical testing with a Benjamini-Hochberg FDR correction cut off of 0.05, ANOVA significant hits were then submitted to a post-hoc test with FDR of 0.05 and the heatmap generated. Cluster analysis was performed to generate the enrichment factor (EF) of specific terms in twelve clusters using the Fisher exact test with a Benjamini-Hochberg FDR threshold of 0.02. All GO/KEGG terms displayed had a p value < 0.0005, N=4.

The first cluster comprised of major histocompatibility complex (MHC) protein binding, toll-like receptor (TLR) signalling, and endocytosis which were increased in J774A.1, RAW 264.7 and BMA3.1A7 cells. The processes enriched in cluster 2 were predominantly associated with cellular homeostatic ion balance and were higher in all cell lines, except for BMA3.1A7 and WT BMDMs. Cluster 3 exhibited enrichment in all non-primary cell lines and was associated with terms related to the innate inflammatory response orchestrated by macrophages, including adenosine deaminase and nitric oxide synthase regulator activity, and MHC binding. Indicative of changes in different pathways that contribute to the eradication of pathogens. Regulation of retroviral genome regulation was observed, which may be lower in J774A.1 cells due to its origin from a murine cancer compared with the other cell lines which were generated using retroviral methods.

Clusters 4, 5, and 6 were elevated in all cell lines. These terms linked to increased cell cycle activity, protein synthesis, metabolism, and biosynthetic process. These findings relate to cell cycle and the increased cell and protein turnover in non-primary cells, which occurs slower in primary cells as they reach senescence (Mathieson et al., 2018).

Cluster 7 referred to terms mainly suppressed in all cell lines. Suppression of podosome assembly, ERK1/2 cascade, regulation of early to late endosome transport, and defence response to bacteria indicates a decreased innate response ability of cell lines.

The terms enriched in cluster 8 were suppressed highly in J774A.1 and BMA3.1A7 cells and all relate strongly to the uptake and clearance of particulate material by macrophages. J774A.1 cells most closely matched BMDMs in cluster 9, which again links closely to phagocytosis.

The terms enriched in cluster 11 were downregulated specifically in BMA3.1A7 cells. The suppression of these terms indicates that BMA3.1A7 cells may possess altered inflammatory signalling and phagocytic capacity.

Finally, in cluster 12, J774A.1 cells were seen to have suppressed activity of the enriched processes. These include the proinflammatory stress-activated protein kinase (SAPK/JNK) signalling cascade and insulin-like growth factor receptor (IGFR) binding.

Overall, the comprehensive comparison between macrophage cell lines and primary macrophages presented in this study offers a broad understanding of their similarities and differences. It sheds light on various crucial processes that may exhibit alterations across different cell types. However, given the extensive size of this dataset, more targeted pairwise comparisons were performed to identify and establish specific differences and proteins that could potentially account for these variations.

### J2-mediated immortalisation drives significant changes in the inflammatory response and phagocytic activity of iBMDMs

Among the identified proteins, 49% show significant differential expression between primary WT BMDMs and WT iBMDMs, with 1696 upregulated and 888 downregulated in WT iBMDMs (**Figure 3**). Gene Set Enrichment Analysis (GSEA) of the entire set of regulated proteins (**Figure 3b and c**) and STRING analysis (**Figure 3d and e**) of the top 50 differentially expressed proteins provided insights into altered protein abundance patterns.

**Figure 3.**
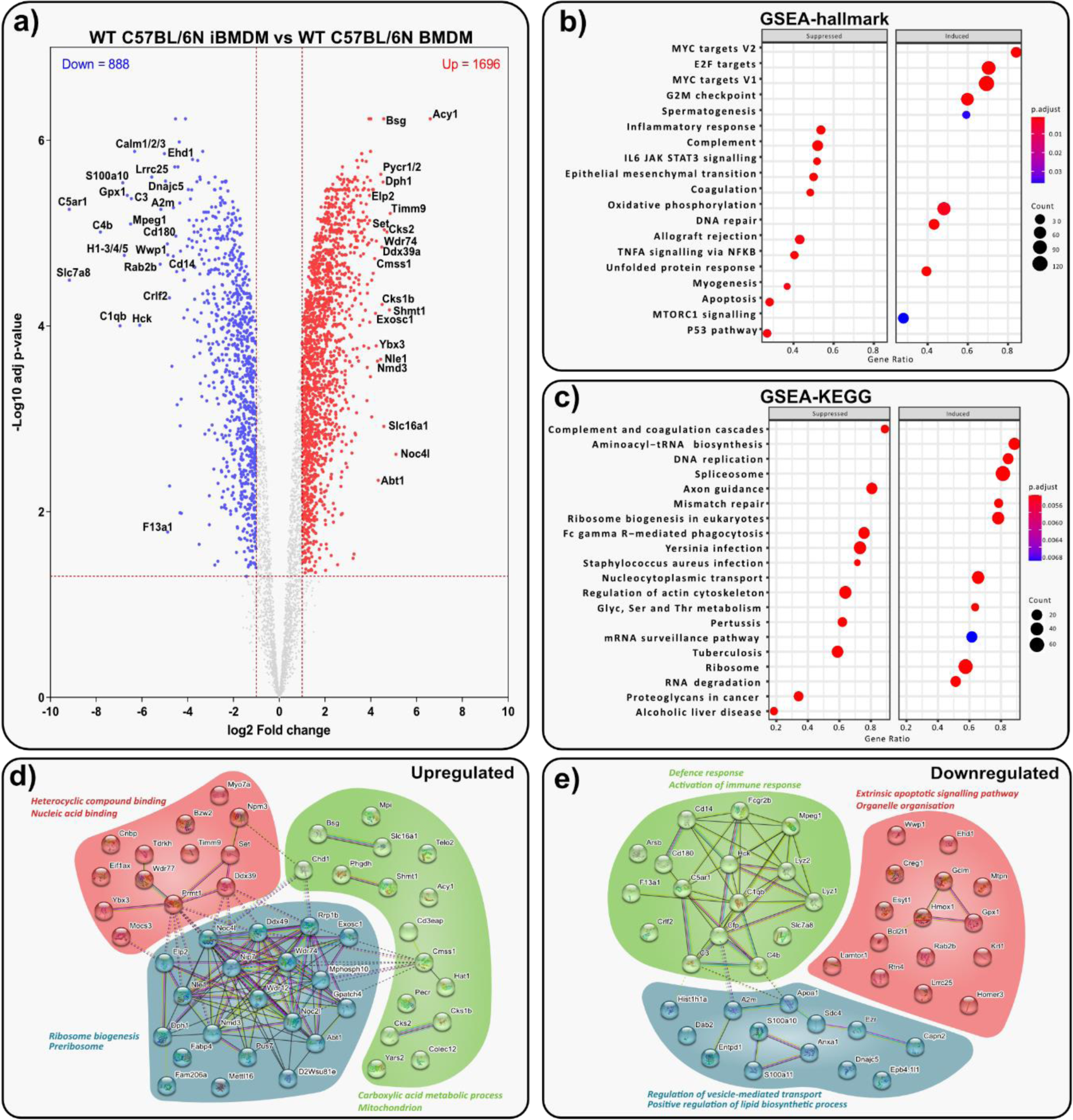
J2-mediated immortalisation drives significant changes in the proteome of BMDMs. **a)** Volcano plot displaying t-test data of WT iBMDMs vs WT BMDMs (log2 fold change > 1 or < -1; > 1.3 –log_10_ (adjusted p-value). Top 20 up- or downregulated proteins are annotated. Analysis of WT iBMDMs vs WT BMDMs shows the top suppressed or induced **b**) hallmark gene sets defined MSigDB and **c)** gene sets defined by KEGG. **d-e)** STRING analysis of the top 50 up and down regulated proteins in WT iBMDMs vs WT BMDMs.

The analyses indicated downregulation of the pro-inflammatory IL-6/JAK/STAT3 signalling axis and a broader inflammatory response in iBMDMs. Conversely, oxidative phosphorylation was induced, suggesting an anti-inflammatory profile compared to WT BMDMs. The complement and coagulation cascades were significantly suppressed, as were Fc-gamma mediated phagocytosis and the bacteriolytic enzymes Lyz1 and Lyz2. STRING analysis revealed upregulation of ribosome biogenesis in iBMDMs, essential for sustained growth.

Overall, iBMDMs exhibited substantial proteomic changes compared to BMDMs, with downregulation of key immune responses and polarization towards an anti-inflammatory phenotype post-immortalization.

### J774A.1 cells are functionally similar to WT BMDMs

J774A.1 cells were revealed to be hierarchically closest to primary macrophages (**Figure 2**). A total of 2465 proteins were shown to be differentially expressed compared to WT BMDMs in **Figure 4a**. However, despite these differences, J774A.1 cells were found to be functionally similar to BMDMs, with few vital processes or pathways affected. Some key proteins were identified as significantly differentially expressed (**Figure 4a**). For example, several proteins pertaining to M2 signalling were significantly downregulated, including GPNMB, MRC1 (CD206), and the tetraspanin CD9. The downregulation of these proteins may indicate a decreased propensity for M2 polarisation in J774A.1 cells, however terms relating to anti-inflammatory M2 processes were not identified in gene set analysis, suggesting that the number of M2 related proteins seen to change was not significant. Two of the main upregulated proteins, aurora kinase B (AURKB) and pyrroline-5-carboxylate reductase 1/2 (PYCR1/2), are oncogenes with increased expression in multiple malignancies and associated with MYC in cancer and therefore link to the MYC target hallmarks shown in **Figure 4b** (Burke et al., 2020; Zhao et al., 2022).

**Figure 4.**
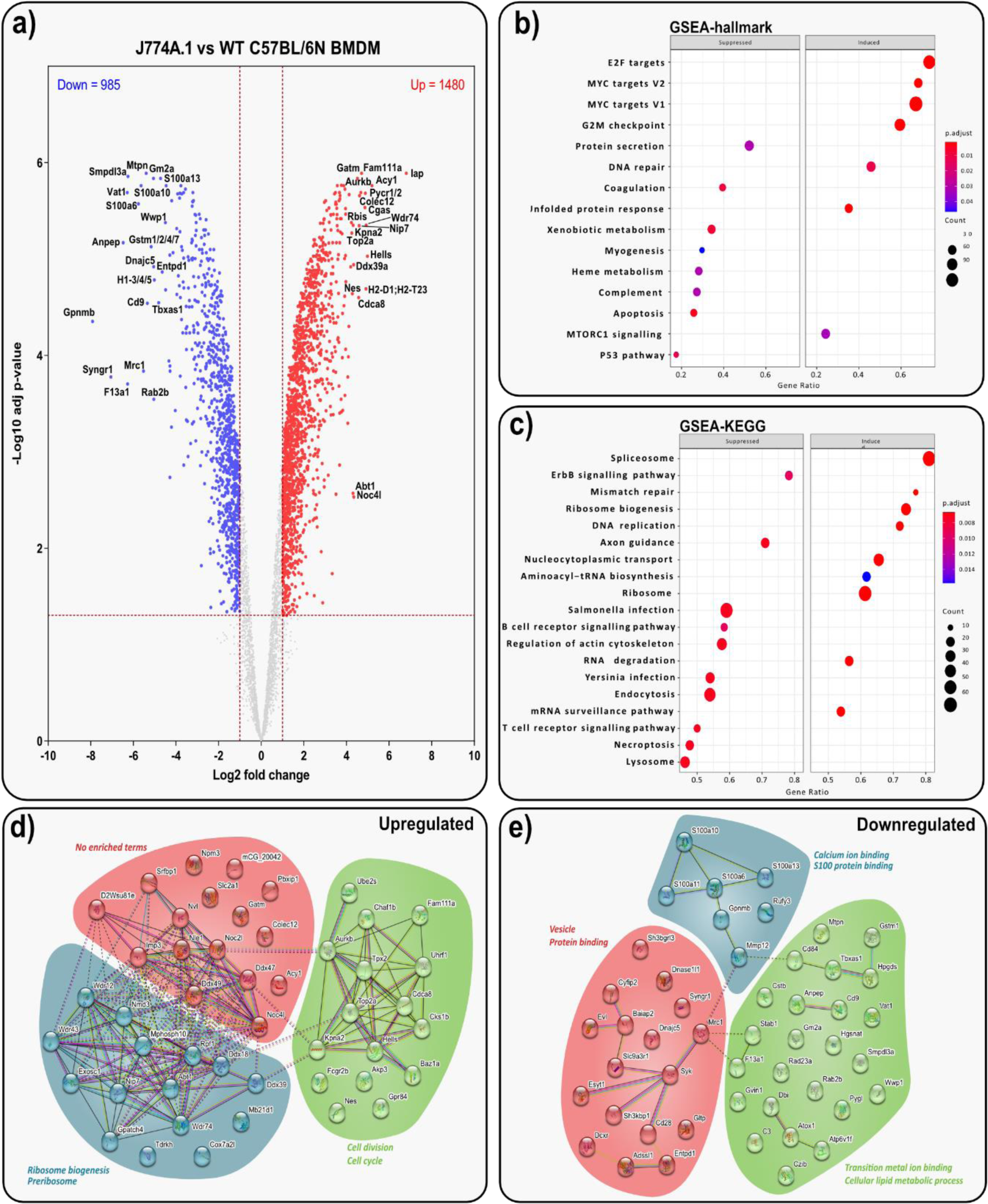
J774A.1 cells are functionally similar to WT BMDMs. **a)** Volcano plot displaying adjusted t-test data of J774A.1 cells *vs* WT BMDMs (log2 fold change > 1 or < -1; > 1.3 –log_10_ (adjusted p-value). Top 20 up- or downregulated proteins are annotated. Analysis of J774A.1 cells *vs* WT BMDMs shows the top suppressed or induced **b**) hallmark gene sets defined MSigDB and **c)** gene sets defined by KEGG. **d-e)** STRING analysis of the top 50 up and down regulated proteins in J774.A1 cells vs WT BMDMs.

Our results indicated that J774A.1 cells might have decreased pathogen killing ability with KEGG terms such as *Salmonella* and *Yersinia* infection suppressed, alongside reduced endocytosis and lysosome activity (**Figure 4c**). This indicates that proteins involved in the binding of specific pathogens and subsequent downstream signalling are downregulated. Among these, MRC1 is one such downregulated protein, linked to the recognition and endocytosis of various microorganisms including *Yersinia pestis* (Azad et al., 2014). The main downregulated interacting proteins were associated with S-100/ICaBP type calcium binding domain, and phospholipase A2 inhibitor activity (**Figure 4e**). In addition, the main upregulated proteins in J774A.1 cells are interconnected (**Figure 4d**) and related to ribosomal and preribosomal function such as the Noc complex and Mpp10 complex, which are known to play significant roles in protein biosynthesis.

Overall, J774A.1 cells hold the closest proteome profile to BMDMs, as evidenced by the low number of macrophage function-related proteins showing significantly differential expression. However, changes in M2 function were seen, with decreased abundance of proteins such as GPNMB, CD206, and CD9. Proinflammatory signalling via S100 proteins was also identified by STRING analysis as downregulated. This again indicates altered inflammatory signalling in immortal cell lines.

### Proteins involved in inflammatory responses are altered in RAW264.7 cells compared to WT BMDMs

RAW264.7 cells exhibit lower similarity to primary WT BMDMs compared to J774A.1 cells (**Figure 2**). Some proteins differentially expressed in J774A.1 cells, such as CD9 downregulation and PYCR upregulation, are also observed here, indicating cancer-driven metabolic reprogramming. Abundance changes of proteins that directly relate to macrophage function were observed (**Figure 5a**). Downregulation of proinflammatory protein MPEG1 and upregulation of insulin-like growth factor 2 receptor (IGF2R) suggest decreased proinflammatory ability. However, upregulation of GLUT1 may enable RAW264.7 cells to acquire an M1 metabolic phenotype more readily. Increased proinflammatory activity is further supported by the upregulation of cellular nucleic acid–binding protein (CNBP) and MHC class I member H2-D1. Pre-ribosomal proteins are also significantly upregulated in RAW264.7 cells (**Figure 5d**).

**Figure 5.**
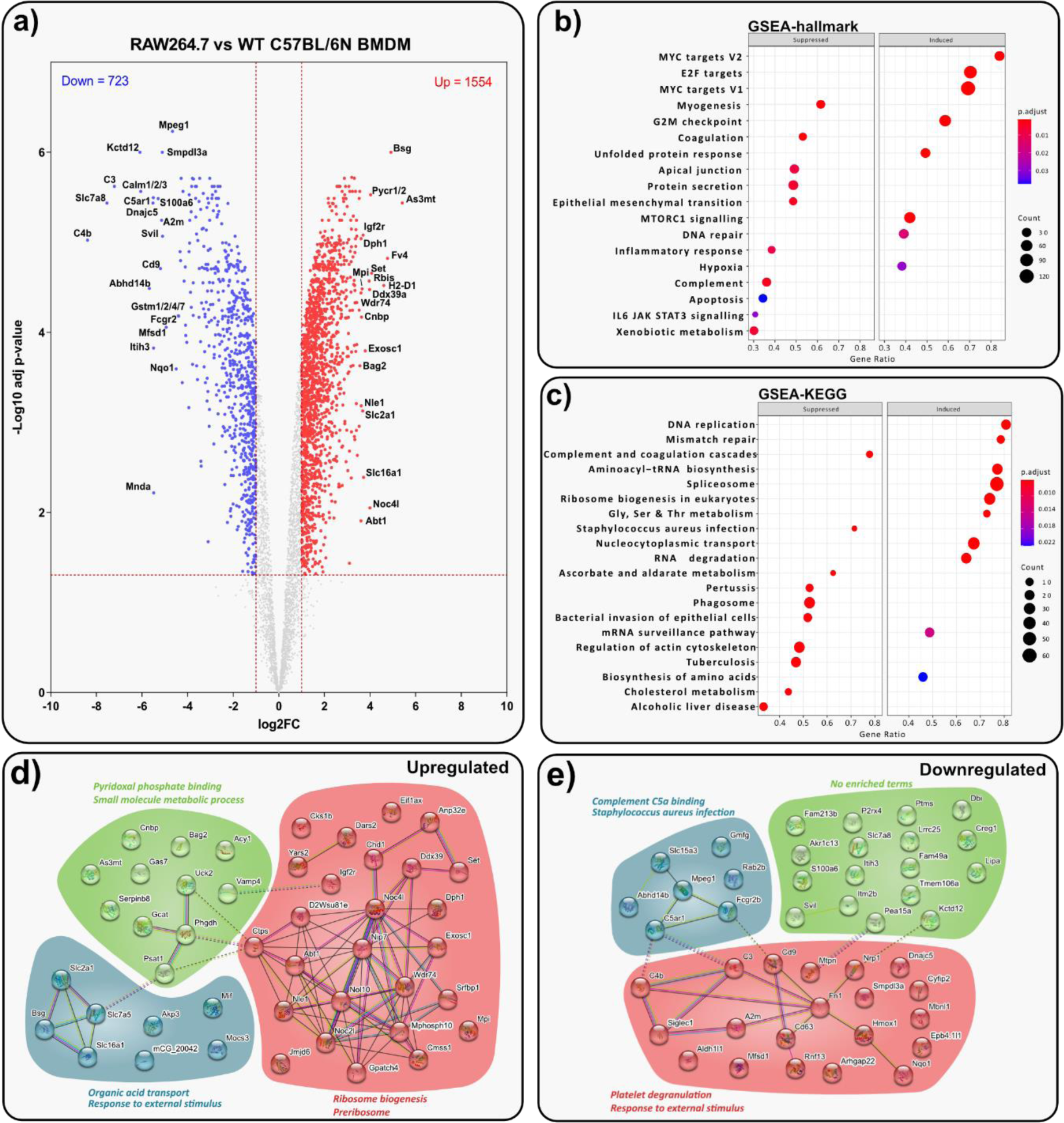
Inflammatory response is altered in RAW264.7 cells compared to WT BMDMs. **a)** Volcano plot displaying t-test data of RAW264.7 cells vs WT BMDMs (log2 fold change > 1 or < -1; > 1.3 –log_10_ (adjusted p-value). Top 20 up- or downregulated proteins are annotated. Analysis of RAW264.7 cells vs WT BMDMs shows the top suppressed or induced **b**) hallmark gene sets defined MSigDB and **c)** gene sets defined by KEGG. **d-e)** STRING analysis of the top 50 up and down regulated proteins in RAW264.7 cells vs WT BMDMs.

GSEA reveals changes in the IL6/JAK/STAT3 signalling axis and inflammatory response hallmarks (**Figure 5b and c**). KEGG terms related to infections and phagosome activity are downregulated in RAW264.7 cells, suggesting a weakened ability to signal in a proinflammatory, host-protective manner. Similar to iBMDMs, complement and coagulation-related proteins (C3, A2m, C4b, C5ar1) were significant downregulated (**Figure 5e**).

### The BMA3.1A7 proteome profile is significantly different to that of WT BMDMs

BMA3.1A7 cells were identified as least similar to WT BMDMs by hierarchical clustering (**Figure 2**), this separation is recapitulated in **Figure 6** which displays a wide range of important differences. Several proteins vital for normal macrophage function were identified as significantly downregulated (**Figure 6a**). These include MPEG1, CD84, TLR-13, LYZ1/2, and CD206. These changes all indicate a decreased ability of BMA3.1A7 cells to recognise and respond to pathogens, overall resulting in lower pro-inflammatory M1 activity and signalling (Abdelaziz et al., 2020; Bayly-Jones et al., 2020; Kolter et al., 2016; Ragland and Criss, 2017; Sintes et al., 2010).

**Figure 6.**
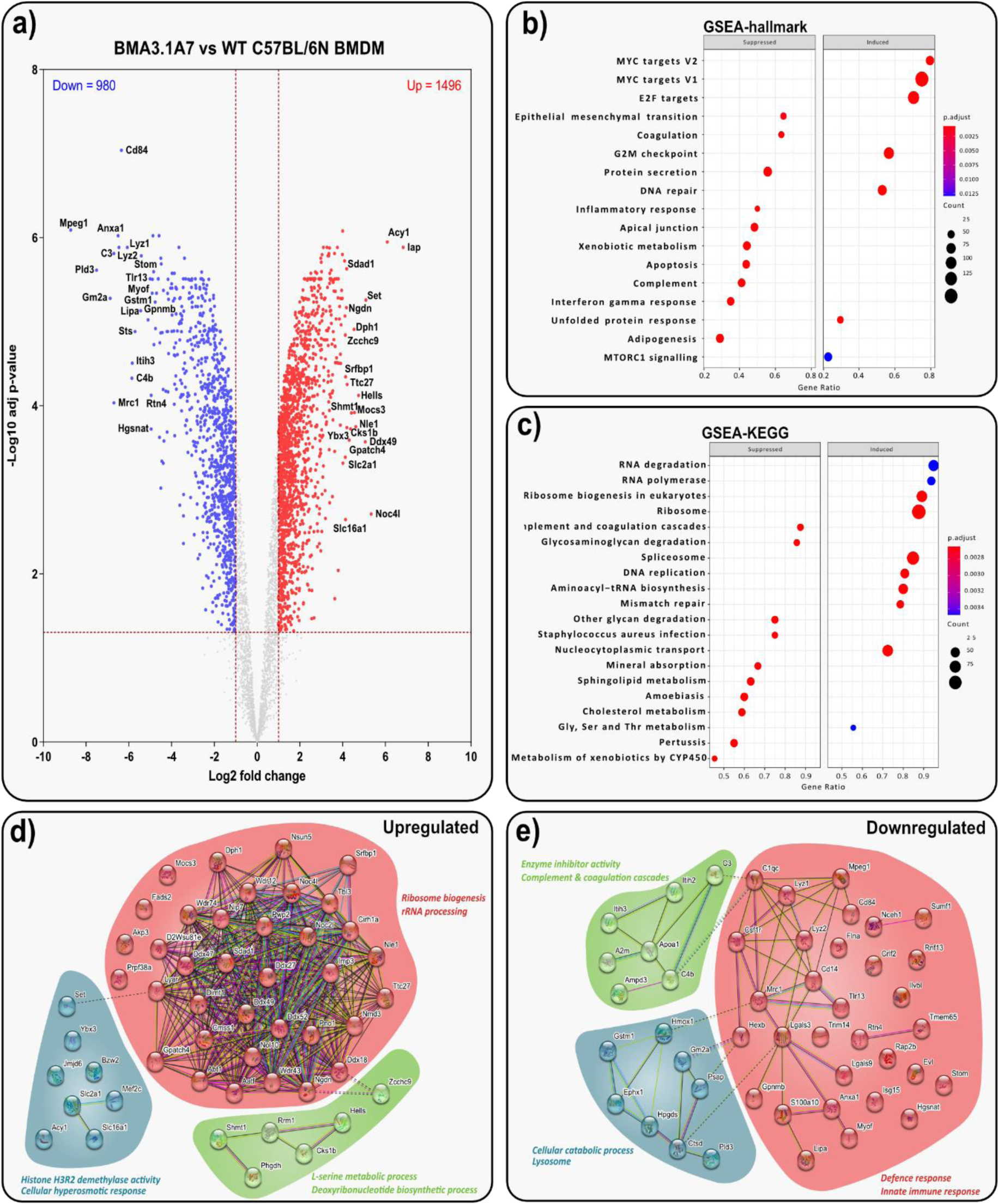
BMA3.1A7 phenotype is significantly different to that of WT BMDMs. **a)** Volcano plot displaying t-test data of BMA3.1A7 cells vs WT BMDMs (log2 fold change > 1 or < -1; > 1.3 –log_10_ (adjusted p-value). Top 20 up- or downregulated proteins are annotated. Analysis of BMA3.1A7 cells vs WT BMDMs shows the top suppressed or induced **b**) hallmark gene sets defined MSigDB and **c)** gene sets defined by KEGG. **d-e)** STRING analysis of the top 50 up and down regulated proteins in BMA3.1A7 cells vs WT BMDMs.

GSEA results show that the inflammatory and IFN-γ responses are suppressed (**Figure 6b**), as well as *Staphylococcus aureus* infection response (**Figure 6c**). Moreover, the downregulation of TLR-13 may contribute to the decreased response to *Staphylococcus aureus* infection as it has been shown to be essential for its recognition in BMDMs (Kolter et al., 2016).

STRING analysis highlighted a highly interconnected cluster of upregulated proteins, containing the highest number of proteins relating to ribosomal and pre-ribosomal process compared to each other cell line (**Figure 6d**). Making up part of this cluster are the DEAD/DEAH box helicase proteins. The other main constituents of the large cluster are WD40 repeat (WDR) containing proteins. The WDR domain is a highly abundant protein interaction domain involved in a variety of processes including the ubiquitin-proteasome system, cell cycle control, regulation of gene expression, and immune responses (Jain and Pandey, 2018; Schapira et al., 2017).

The downregulated proteins identified by STRING analysis again link strongly to macrophage function and phenotype (**Figure 6e**). Galectin-3 (Lgals3) and -9 (Lgals9) are both linked to inflammatory signalling. Proteins linked to the classical pathway of complement activation were also seen to be downregulated again. Importantly, BMA3.1A7 cells were the only cells to have the terms ‘bone marrow-derived macrophage’ and ‘innate immunity’ highlighted in the downregulated STRING analysis.

Taken together, these results suggest that the BMA3.1A7 cell line exhibits the lowest degree of macrophage-like characteristics, as evidenced by the enriched terms identified in the downregulated STRING analysis. Furthermore, most of the notable alterations observed were associated with significant changes in pro-inflammatory or pattern recognition receptor signalling pathways, implying that this cell line may be less suitable for studies related to innate immune response.

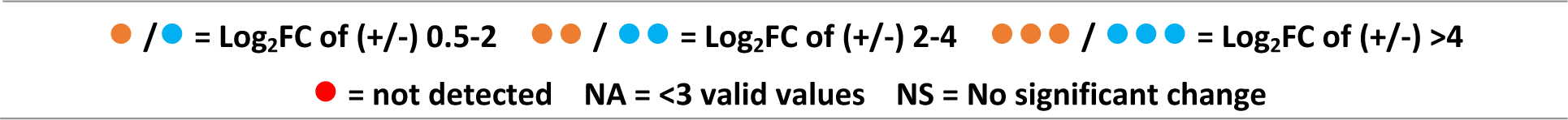

### Macrophage receptor expression is significantly altered in macrophage cell lines

Macrophages employ a large catalogue of intracellular and surface receptors to act as sensors allowing identification of their surroundings cells, the state of the tissue, various metabolites, lipoproteins, antibodies, complement, and pathogens (Ley et al., 2016). These receptors act in tandem to regulate the protective function of macrophages. Therefore, it is essential that these receptors are expressed in cell lines at similar levels to primary macrophages. Macrophage receptors identified in this dataset were extracted along with their fold change in comparison to WT BMDMs (**Table 1**), providing potential insights into choosing the right cell line for experiments.

**Table 1.**
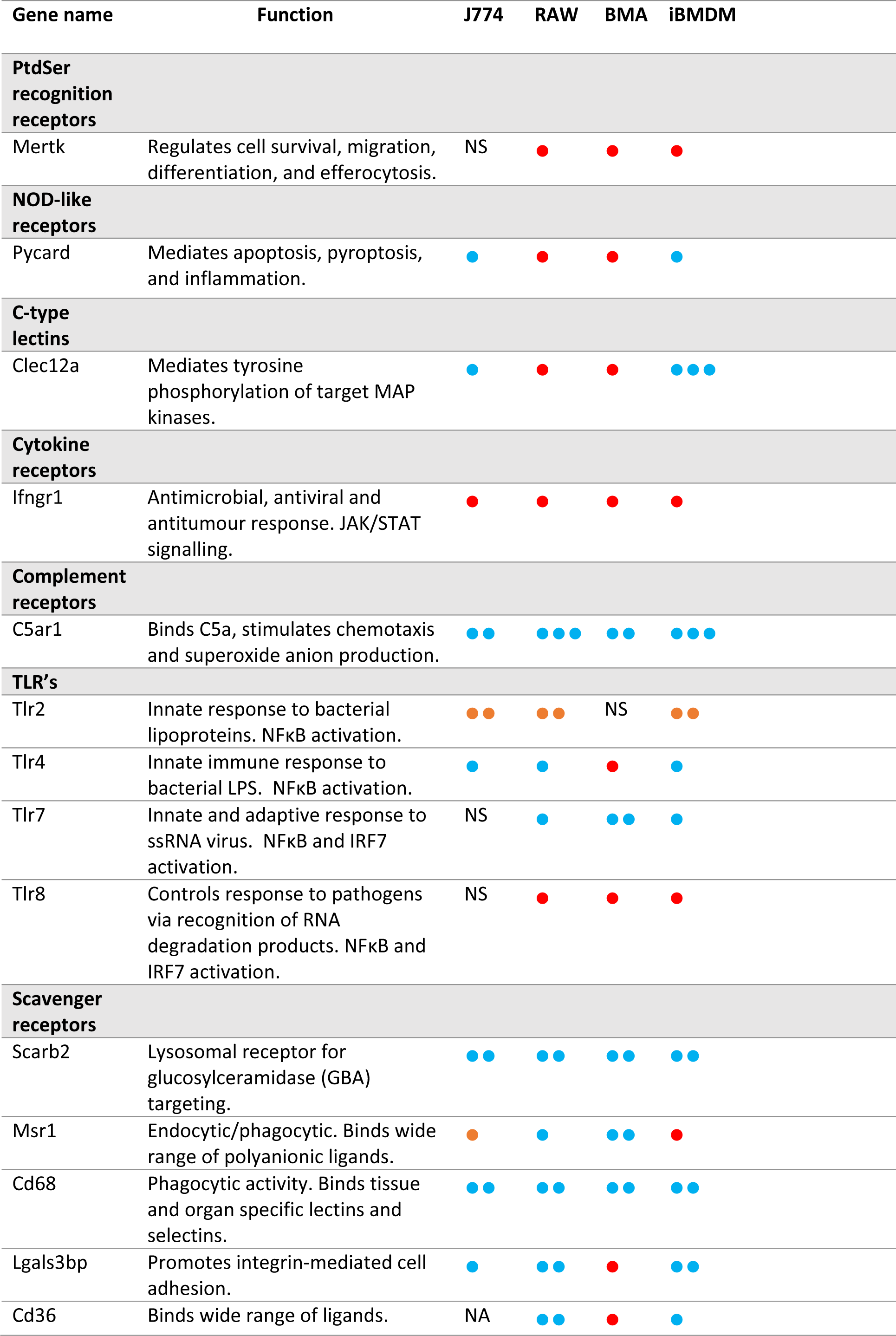
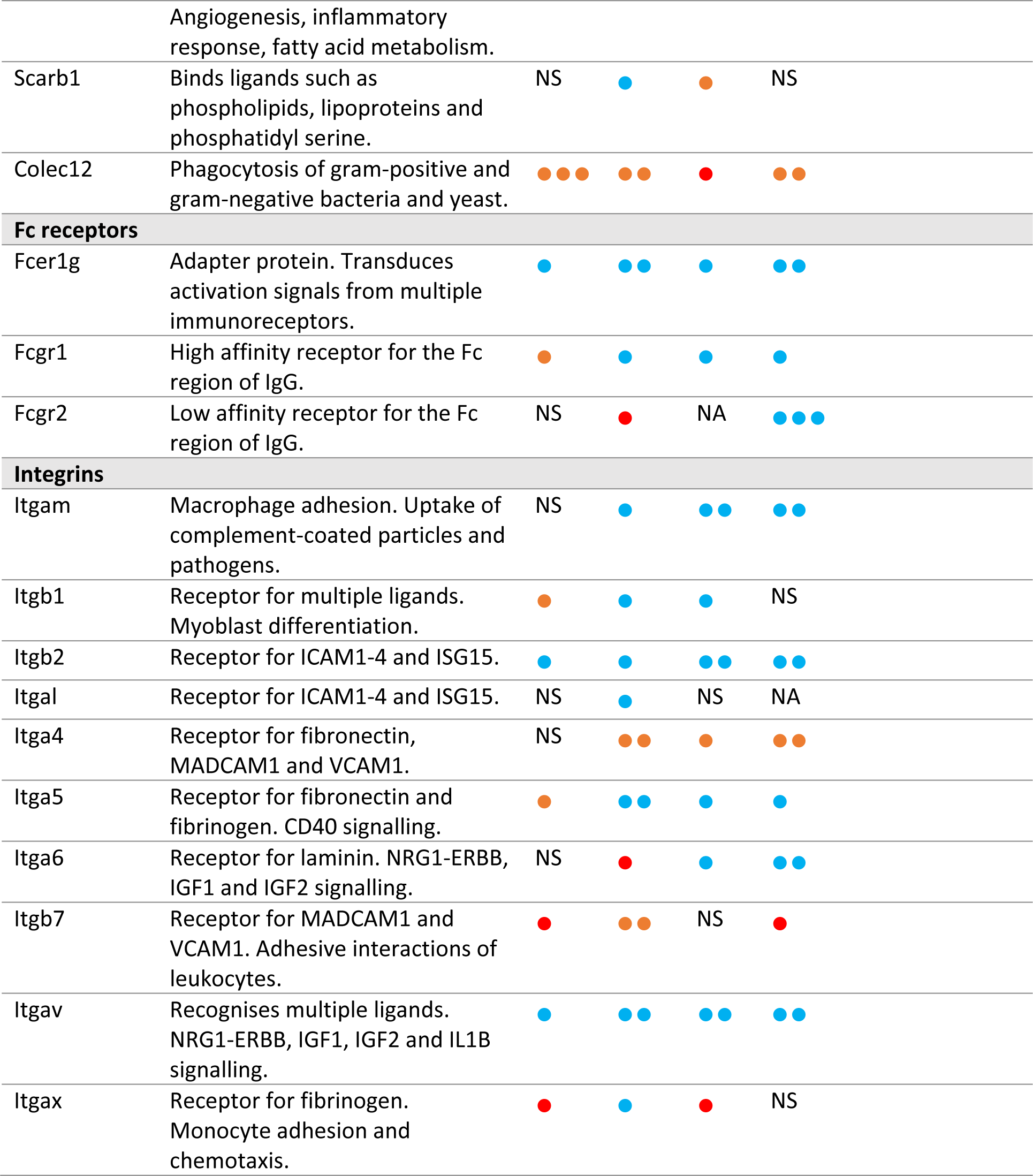
Macrophage receptor expression in macrophage cell lines in comparison to WT BMDMs. Protein name and functions are indicated. Fold changes are presented in coloured circles as indicated in the legend. Log2 fold change (Log2FC) was calculated for each cell line in comparison to WT BMDMs. Unless otherwise stated, all values are significant at <0.05 adjusted p-value. N=4.

### Proteins crucial to macrophage function were unidentified in certain cell lines

Both global analysis and individual comparisons of proteome differences between cell lines and WT BMDMs gave valuable insight into the suitability of cell lines for the immunology and microbiology infection fields. However, these comparisons only considered proteins which were identified in at least three out of four biological replicates for each group. This meant that proteins completely unidentified in one cell type, but expressed in another, were not considered, as the data was not imputed. Such proteins might be of high importance when analysing experimental results or considering which cell line to use for specific investigations. We performed heatmap analysis of proteins identified in all four biological replicates of WT BMDMs but not in any replicate of at least one of the cell lines under investigation. These proteins were grouped into three hierarchical clusters labelled as (a), (b), and (c) (**Figure 7**).

**Figure 7.**
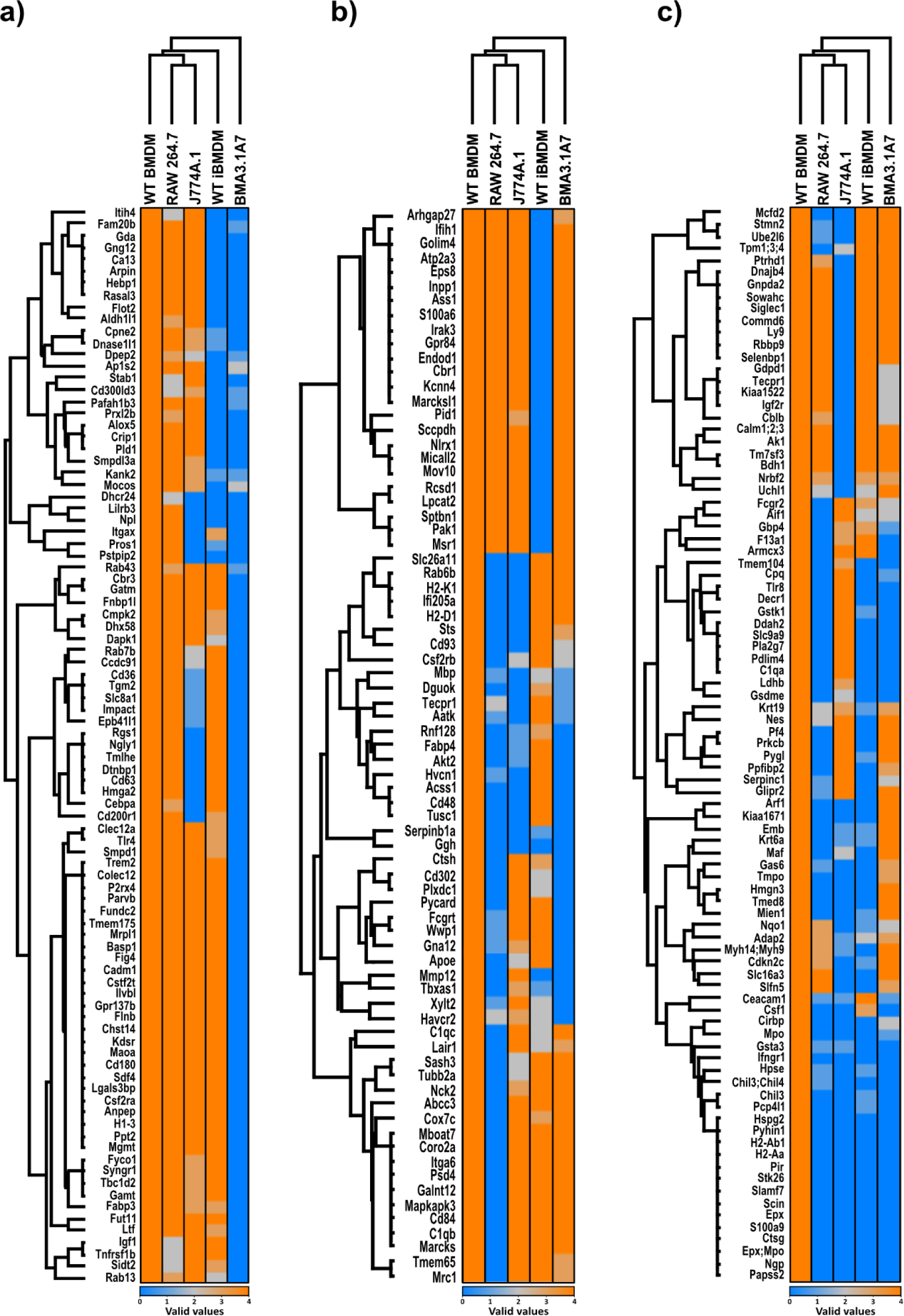
Cell lines lack key proteins that are present in WT BMDMs. Presence/absence was determined using the number of valid values identified by mass spectrometry for each protein. Presence defined as three or more valid values (orange) and absence defined as zero valid values (blue). Only proteins displaying four valid values in WT BMDMs were included. Hierarchical clustering was performed in Perseus with the resulting heatmap separated into three main clusters (a, b, and c). N=4.

Cluster (a) of the heatmap displays proteins that were mainly not detected in BMA3.1A7 cells. GSEA of this set of proteins revealed that, in general, this cluster was related to the response to external stimulus, innate immune response-activating signalling pathway, and defence response (**Figure 7a**). In particular, scavenger receptor activity was highlighted in this cluster, with stabilin-1 (STAB1), CD36, and collectin-12 (COLEC12) all absent in BMA3.1A7 cells. Other proteins important for host defence were also not identified, such as transglutaminase 2 (TGM2), triggering receptor expressed on myeloid cells 2 (TREM2), and TLR-4.

Unlike in cluster (a) the proteins displayed in cluster (b) were not identified in various cell lines (**Figure 7b**). However, the proteins here were again linked to immune and defence response. Several proteins relating to FcγR-mediated phagocytosis were not identified in various cell lines, potentially giving rise to differences in the uptake of opsonised particles. Myristoylated alanine rich c-kinase substrate (MARCKS) was absent in RAW264.7 cells as was MARCKS-like protein 1 (MARCKSL1) in WT iBMDMs. These proteins affect processes such as cytoskeletal rearrangement, vesicular trafficking, and phagocytosis (El Amri et al., 2018). AKT2, a serine-threonine kinase, was not identified in BMA3.1A7, RAW264.7 and J774A.1 cells. Deficiency of this kinase shifts polarisation towards an M2-like state, negatively regulates TLR-4 signalling, and impedes the uptake of opsonised beads (Vergadi et al., 2017). Inflammasome function may be affected in RAW264.7 and BMA3.1A7 cells by the lack of PYCARD, an adaptor protein essential for the formation of the AIM2 and NLRP3 inflammasome (Fang et al., 2019). CD93, not identified in RAW264.7 and J774A.1 cells, and CD48, not detected in all three common cell lines, might also impact host-protection against bacteria (McArdel et al., 2016; Nativel et al., 2019). Whilst varied in their functions, the proteins identified in this cluster are strongly related to inflammatory signalling and pathogen clearance. This indicates that cell lines might display varying responses to certain stimuli depending on which signalling pathway members are present or absent.

Cluster (c) contains proteins undetected in most of the cell lines and relate to coagulation, defence response to another organism, wound healing, immune response, and regulation of cytokine production (**Figure 7c**). IGF2R was not identified in J774A.1 cells. This protein, known to regulate anti-inflammatory metabolic reprogramming, was previously shown to be significantly upregulated in RAW264.7 cells (**Figure 5**). Therefore, M2 polarisation and signalling may differ between these two cell lines. S100a9 normally functions in complex with S100a8 in response to infection by inducing pro-inflammatory cytokines, reactive oxygen species and nitric oxide. It also mediates rearrangement of the cytoskeleton and is therefore implicated in phagocytosis (Wang et al., 2018). H2-AA and H2-AB1, both class II histocompatibility antigens, are important for the presentation of antigens to CD4+ T cells by MHC class II (Stables et al., 2011). Therefore, part of the adaptive immune response is impaired in these cells. Finally, and importantly, Interferon-γ receptor 1 (IFNGR1) was not identified in any cell line.

## Discussion

The information provided here highlights the importance of cell line characterisation in comparison to the gold standard primary C57BL/6 BMDM model. Such characterisations enable selection of suitable models and insight into why differences in the response of cell lines to stimuli may differ.

Induction of common functions and pathways across all cell lines was highlighted, such as *Myc* activity, G2-M checkpoint, E2F gene transcription and ribosome activity. This is indicative of the cellular changes needed to support continuous proliferation. MYC is a key transcription factor known to contribute to the development of several cancers. Specifically, *Myc* V1 targets include genes involved in cell cycle regulation, DNA replication, and protein synthesis, while V2 targets include genes involved in apoptosis and immune response. V1 targets are upregulated by *Myc*, whereas V2 targets are downregulated (H. Chen et al., 2018; Dang, 2012). E2F targets and G2/M checkpoint also link to the immortalisation process and aberrant cell cycle progression (Oshi et al., 2020b, 2020a).

Complement and coagulation cascade related proteins were also seen to be downregulated in each cell line. These cascades form part of the first line of innate defence against invading pathogens in the circulation, with both systems relying on complex enzymatic signalling cascades for efficient activity. The same changes were seen in a proteome comparison of the hepatoma cell line Hepa1-6 with primary hepatocytes (Pan et al., 2009). Therefore, it is likely that loss of tissue context means that it is no longer necessary for the cell lines to produce these proteins. However, besides these common differences, each cell line displayed varied changes relating to normal macrophage function and innate immunity.

The J2 immortalisation process induced changes mainly related to macrophage polarisation. The proinflammatory JAK/STAT pathway was suppressed in iBMDMs, this pathway is linked with increased tumour cell proliferation and dampened antitumor immune response. In the tumour microenvironment, the IL-6/JAK/STAT3 axis is stimulated by release of IL-6 from cells such as TAMs (Johnson et al., 2018). Furthermore, proteins relating to oxidative phosphorylation were significantly induced. M2 polarised macrophages rely on oxidative phosphorylation for energy metabolism, whereas M1 macrophages rely mainly on glycolysis (Viola et al., 2019). The reduction in complement activity seen in these cells, specifically activity of C5a and C3a, may also be partially responsible for the suppression of the IL6/JAK/STAT3 cascade, as C5a and C3a are known to stimulate IL6 release in Kupffer cells (Oikonomopoulou et al., 2012). Further decreased ability to respond to and clear pathogens was indicated by the downregulation of Fc-γ mediated phagocytosis and Lyz1/2. Fcγ receptors on macrophages mediate the uptake and killing of foreign material such as antibody-coated bacteria, viruses, and parasites as well as host cells that express viral or tumour antigens (Fitzer-Attas et al., 2000). Whilst Lyz1/2 are important enzymes capable of hydrolysing bacterial cell wall peptidoglycans (Ragland and Criss, 2017).

J774A.1 cells were determined to be the most similar to WT C57BL/6 BMDMs, despite being a BALB/c model. However, the cell line harboured some important changes. Knockdown of the most downregulated protein in J774A.1 cells, GPNMB, in RAW264.7 cells resulted in increased levels of pro-inflammatory cytokines, whilst overexpression dampened the M1 response after IFN-γ/LPS stimulation (Saade et al., 2021). Also downregulated, MRC1, plays multiple roles in macrophage activity, including the recognition and endocytic clearance of both endogenous and pathogen-related ligands, antigen processing and presentation, and TLR-2 signalling (van der Zande et al., 2021). Furthermore, linking to the M2 phenotype, ablation of the receptor increases levels of pro-inflammatory proteins. Similarly, CD9 has been shown to negatively regulate LPS induced macrophage activation (Brosseau et al., 2018; Orecchioni et al., 2019). Downregulation of these proteins suggests a shift towards M1 activity. Several S100 proteins were also downregulated. The S100 proteins hold a range of functions but have importantly been linked with the regulation of macrophage inflammation. S100a6 interacts with receptor for advanced glycation end products (RAGE) to mediate cell survival and apoptosis and activates proinflammatory JNK. S100A10 has been shown to mediate migration of macrophages towards tumour sites and contribute to MAPK and NF-κB induction of inflammatory cytokines. S100A11 is an alarmin, an endogenous chemotactic and immune activating protein, secreted in response to *Toxoplasma gondii.* S100A13 also further contributes to cytokine and antimicrobial agent release (Singh and Ali, 2022; Xia et al., 2018).

Changes seen in RAW264.7 cells were more varied. MPEG1 was downregulated, this is normally induced in response to proinflammatory stimuli, with its loss linked to increased susceptibility to bacterial infection (Bayly-Jones et al., 2020). Additionally suggesting decreased proinflammatory ability, insulin-like growth factor 2 receptor (IGF2R) was upregulated. Activation of IGF2R reprograms cellular metabolism towards oxidative phosphorylation and thus an anti-inflammatory phenotype (Wang et al., 2020). However, also implicated in metabolic reprogramming, SLC2A1 (GLUT1) was upregulated. This is the main rate-limiting glucose transporter found on M1 macrophages. M1 macrophages favour glucose as an energy substrate, therefore RAW264.7 cells may more readily acquire an M1 metabolic phenotype (Freemerman et al., 2014). Further supporting increased proinflammatory activity in RAW264.7 cells, CNBP is known to regulate a specific gene signature in M1 macrophages, by controlling the activity of c-Rel and its translocation to the nucleus. This activity specifically controls the induction of IL-12β expression and therefore impacts the T helper type 1 and IFNγ-mediated immune response against a range of pathogens (Y. Chen et al., 2018).

BMA3.1A7 cells harboured the greatest number of significant changes in proteins linking strongly to normal macrophage function, with many downregulated or not detected. Firstly, MPEG1 was downregulated 256-fold, potentially increasing susceptibility to bacterial infection by decreasing the pro-inflammatory response (Bayly-Jones et al., 2020). CD84 has been implicated in the LPS response by rapidly increasing phosphorylation of ERK-1/2, p38, and JNK-1/2; and by activating NF-κB (Sintes et al., 2010). Downregulation may therefore hinder the ability to fully respond to LPS. TLR-13 recognises a conserved motif present in 23S ribosomal RNA from both gram-positive and gram-negative bacteria (Li and Chen, 2012), as well as vesicular stomatitis virus (Shi et al., 2011). Group B streptococcus was used to demonstrate that upon ligand binding, TLR-13 mediates TNF-α and nitric oxide induction (Signorino et al., 2014). Lyz 1 and 2 were again seen to be significantly downregulated, limiting protection against invading pathogens (Ragland and Criss, 2017). CD206, commonly referred to as an M2 macrophage marker that recognizes various PAMPs was also downregulated. (Abdelaziz et al., 2020). Galectin-3 inhibition can reduce inflammatory response and the expression of markers such as TNF-α and IL-1β (Lu et al., 2020). Whereas galectin-9 can stimulate monocyte differentiation towards an M2 phenotype (Enninga et al., 2016). Further to significantly downregulated proteins, several key proteins were not identified at all in BMA3.1A7 cells. STAB1 acts as an endocytic receptor for self-ligands such as acLDL and as a phagocytic receptor for apoptotic bodies (Kzhyshkowska, 2010). Similarly, CD36 has high affinity for specific phospholipid moieties found on oxLDL and apoptotic cells and also binds glycated proteins, amyloid β, LTA from bacteria, and glycans from fungi. Recognition of these DAMPs or PAMPs results in a pro-inflammatory response mediated by NF-κB (Chen et al., 2022). COLEC12 may be responsible for clearance of amyloid β (Kelley et al., 2014). TGM2 is known to promote efferocytosis, with ablation of the enzyme reducing levels of efferocytic receptors CD14 and MSR1, and promoting pro-inflammatory signalling (Eligini et al., 2016). TREM2 is a phagocytic receptor for bacteria and amyloid β (Li and Zhang, 2018). Finally, TLR-4 is the major toll-like receptor for LPS and other exogenous ligands, implicating the receptor in host defence against gram-positive and gram-negative bacteria, viruses, and fungi. TLR-4 also recognises endogenous ligands such as heat-shock proteins, hyaluronan, and amyloid β. The loss of these proteins in BMA3.1A7 cells, amongst the many others outlined in **Figure 7**, indicates that this cell line has an impaired ability to recognise exogenous PAMPs and endogenous DAMPs. Therefore, this cell line may not be suitable for experiments focussed on phagocytosis, efferocytosis or infection models such as *Staphylococcus aureus* infection.

Investigation of receptor expression highlighted that J774A.1 cells hold the most similar receptor profile to WT BMDMs. Mer tyrosine kinase (MERTK) was only identified in J774A.1 cells, this receptor plays an important role in the regulation of cytokine secretion and apoptotic cell clearance. Furthermore, mice deficient in MERTK are hypersensitive to LPS-induced endotoxic shock, indicating that the receptor has a role in the dampening the immune response in macrophages (Anwar et al., 2009). J774A.1 cells therefore may be a more suitable model for exploring the effect of LPS treatment on macrophage function. Expression of TLR-7 and TLR-8 was also most similar to WT BMDMs in J774A.1 cells, both receptors recognize single-stranded RNA and induce an inflammatory response (Eng et al., 2018). This means that these cells may give a more representative response to single-stranded RNA viruses that other models. FCGR2 expression is also normal in J774A.1 cells compared to RAW264.7 cells, indicating that J774A.1 cells may be a better model for investigating the role of macrophages in the phagocytosis and killing of opsonised pathogens.

One of the more puzzling findings was the lack of identification of IFNGR1 in all cell lines. IFN-γ signals by first binding to its dimeric receptor consisting of IFNGR1 and IFNGR2. This activates the JAK-STAT signalling pathway and interferon-stimulated gene (ISG) production. ISGs encode a wide range of products that hold vital immune effector functions such as antigen-presentation, chemokine activity, and phagocytosis. Therefore IFN-γ is one of the most important endogenous inflammatory and immune mediators, vital for host defence against intracellular pathogens (Hu and Ivashkiv, 2009; Ivashkiv, 2018). Lack of identification of this vital receptor across all cell lines was unexpected as each cell line has previously been reported to respond to IFN-γ stimulation (Bilkei-Gorzo et al., 2022; Elizabeth et al., 2016; Gao et al., 2016; Raso et al., 2002).

The caveat to the proteomic comparisons performed here is that all comparisons were made to C57BL/6 BMDMs. Both J774A.1 and RAW264.7 cells were obtained from BALB/c mice, with J774A.1 cells generated from ascites of reticulum cell sarcoma (Ralph et al., 1976) and RAW264.7 cells produced by immortalisation with Abelson murine leukaemia virus (Raschke et al., 1978). Therefore, differences at the proteome level may be influenced by the difference in strain. However, macrophage investigations throughout the literature use either BMDMs from C57BL/6 mice or J774A.1 and RAW264.7 cells. Therefore, these comparisons remain valuable. Further differences may also be a result of the method used to generate the cell lines. The J774A.1 cell line displayed the least amount of change relevant to macrophage function and is the only line to not be generated using a virus. This potentially indicates that tumour derived macrophage cell lines are more similar to primary cells than cell lines generated using oncogenic viruses.

It appears that the process of J2-immortalisation drove the unexpected loss of MSR1, as confirmed by mass spectrometry, flow cytometry and RT-qPCR. This has not been noted in any previous literature. The molecular reasoning behind the loss of MSR1 is unclear, however as RT-qPCR analysis indicated diminished gene transcription, it is likely to be a result of genetic or epigenetic interference. One potential mechanism of this is gene disruption due to transgene insertion. As the J2 virus is a gamma-retrovirus, it may have an increased propensity for insertion near promotor regions (Baum et al., 2004). Alternatively, the J2 virus may have induced CpG methylation near the transcription start site or promotor regions of *Msr1*, which could silence the gene. The same mechanism may be responsible for the silencing of other genes discussed here.

The dataset presented here (**Table S1 and S2**) will be of valuable use to the macrophage biology community. Differential expression of various proteins may explain experimental differences obtained when comparing primary models to cell line models. Furthermore, the proteome profiles generated can guide the selection of optimal models for use in experiments covering areas such as phagocytosis, polarisation, and infection.

## Materials and methods

### Cell culture

J774A.1 (TIB-67), RAW264.7 (TIB-71), AMJ2-C11 (CRL-2456), and L929 (CCL-1) cell lines were purchased from the American Type Culture Collection (ATCC, Manassas, VA, USA). The BMA3.1A7 (accession: CVCL_IW58) was kindly provided by Kenneth Rock (Dana Farber Centre, Boston, US). Cells were maintained in medium at 37 °C in a humidified 5% CO_2_ atmosphere. ATCC routinely performs cell line authentication, using short tandem repeat profiling as a procedure. Cell experimentation was always performed within a period not exceeding two months after resuscitation and cells were regularly tested for mycoplasma.

### Mice

Wild-type C57BL/6J mice were obtained from Charles River. Newcastle University ethical committee approved animal work and manipulation was performed under UK Home Office project licence.

### Generation of murine bone marrow derived macrophages (BMDMs)

Bone marrow cells were collected femurs and tibiae of 8 to 12-week-old WT and *Msr1^-/-^* C57BL/6 mice. Collected cells were treated with red blood cell lysis buffer (155 mM NH_4_CI, 12 mM NaHCO_3_, 0.1 mM EDTA) and plated on untreated 10 cm cell culture dishes (BD Biosciences) in IMDM (Gibco) containing 10% heat inactivated FBS, 100 units/ml penicillin/streptomycin (Gibco) and 15% L929 conditioned supplement. After 24 h, the cells in supernatant were transferred to untreated 10 cm Petri dishes (BD Biosciences) for 7 days for the differentiation into bone marrow-derived macrophages (BMDMs) (Heap et al., 2021).

### Immortalisation of murine bone marrow derived macrophages

The AMJ2-C11 cell line was used as the source of J2 virus needed for the immortalisation of the target cell lines. AMJ2-C11 cells were cultured in complete DMEM (AMJ2-C11) until 80% confluent. Then, 75% of these cells were seeded using bone marrow growth medium and incubated for 24 hours. The J2 virus-containing conditioned media was then decanted and clarified by centrifugation at 200 xg for 5 min followed by filtration through a 0.45 µm filter. Aliquots were either used immediately to avoid loss of viral titre or stored at -20°C.

Murine bone marrow derived macrophages were generated as previously described and transduced twice with J2 virus as follows. Five days after harvest bone marrow growth media was replaced with 50% J2-conditioned media and 50% bone marrow growth media for 24 hours, keeping a control dish of cells for comparison. After 24 hours, J2-conditioned media was removed, and the cells allowed to recover for a further 24 hours in fresh media. The first transduction step was then repeated for a final 24 hours. Cells were routinely monitored for growth or death compared to the control and passaged accordingly. One week after transduction the concentration of L929 conditioned medium in the bone marrow growth media was lowered from 20% to 10% with the aim of reducing this concentration to 0% over the next 2-3 months. This process is likely being carried out to wean the cells off the dependency on L929 conditioned medium. If the immortalised BMDMs (iBMDMs) ceased to grow after a reduction in L929 concentration, the concentration was returned to its previous level and the cells allowed to recover before attempting to lower the concentration again.

### Reverse transcription-quantitative polymerase chain reaction (RT-qPCR)

RNA extraction was performed using Qiagen RNeasy mini columns and QiaShredder for homogenisation kit (Qiagen) following manufacturer’s instructions. RNA was quantified using a NanoDrop spectrophotometer. Reverse transcription was performed with 1 µg of template RNA using the Qiagen QuantiTect reverse transcription kit (Qiagen) in a Bio-Rad T100TM Thermal Cycler (Bio-Rad Laboratories) to obtain complementary DNA (cDNA). cDNA was diluted to 5 ng/µl in RNase-free water and 1 µl used for real-time qPCR with Qiagen QuantiNova SYBR Green Master Mix (Qiagen) in a StepOnePlus Real-Time PCR System (Applied Biosystems) according to the manufacturer’s instructions. Each sample was run in triplicate and the 2-ΔΔCt method used to calculate the relative expression of genes (Table 2.5). Expression values of *Gapdh* and *Tbp* genes were used for normalisation.

### Proteomics sample preparation

Whole cell protein lysate was prepared by sonication in SDS lysis buffer (5% SDS, 50 mM TEAB, pH 8.5 in HPLC water supplemented with cOmplete, Protease Inhibitor Cocktail (Sigma)). Protein concentration was quantified using the Pierce BCA Protein Assay (Thermo Fisher Scientific) according to manufacturer’s instructions and measured at 562 nm using the SpectraMaxTM iD3 microplate reader (Molecular Devices). 25 µg of each sample was reduced by addition of tris (2-carboxyethyl) phosphine (TCEP, Pierce) to a final concentration of 10 mM for 30 min at 37°C, and subsequently alkylated with iodoacetamide (IAA) to a final concentration of 10 mM for 30 min at room temperature in the dark. Samples were acidified by addition of 2.5 μl of 12% phosphoric acid and diluted with 165 μl of S-trap binding buffer (90% MeOH, 100 mM TEAB, pH 7.1). The acidified samples were then loaded onto the S-trap spin column (ProtiFi, Huntington NY, USA) and centrifuged at 4,000 xg for 1 min. Each S-trap mini-spin column was washed with 150 µl S-trap binding buffer and centrifuged at 4,000 xg for 1 min, this was repeated five times with rotation of the column between each wash. For protein digestion TEAB (50 mM, pH 8.0) containing sequencing-grade trypsin (1:10 ratio of trypsin:protein) was added to each sample, followed by incubation for 2 hours at 47°C using an unmoving thermomixer (Eppendorf). Peptides were eluted with 40 µl TEAB (50 mM, pH 8.0) and centrifugation at 1,000 xg for 1 min. Elution steps were repeated using 40 µl formic acid (0.2%) and finally 35 µl formic acid (0.2%), acetonitrile (50%). Eluates were combined and dried down using a speed-vac before storage at −80°C.

### High performance liquid chromatography

Dried peptide samples were resuspended in MS sample buffer (Acetonitrile (2% v/v), trifluoroacetic acid (0.1% v/v) in HPLC-grade water) to a concentration of 500 ng/μl and placed into glass autosampler vials. Peptide samples were injected on a Dionex Ultimate 3000 RSLC (Thermo Fisher Scientific), connected to an Orbitrap QE HF mass spectrometer (Thermo Scientific), using a PepMap 100 C18 LC trap column (300 µm ID x 5 mm, 5 µm, 100 Å). This was followed by separation on an EASY-Spray column (50 cm x 75 µm ID, PepMap C18, 2 µm, 100 Å) (Thermo Fisher Scientific). Peptides were separated using a linear gradient of 3 − 35% Buffer B (Acetonitrile (80% v/v), Formic acid (0.1% v/v) in HPLC-grade water; Buffer A being Formic acid (0.1% v/v) in HPLC-grade water) over 120 min for whole cell samples, followed by a step from 35 − 90% Buffer B in 30 sec at 250 nL/min and held at 90% for 4 min. The gradient was then decreased to 3% Buffer B in 0.5 min at 250 nL/min for 20 min. Column temperature was controlled at 45°C.

### Data-independent acquisition LC-MS/MS analysis

Whole cell proteomes were analysed using DIA on an Orbitrap QE HF mass spectrometer. The QE HF mass spectrometer was operated in positive ion, DIA mode. Full MS scan spectra were acquired in the m/z range of 400-1300 with an automatic gain control target of 1e6 and a maximum injection time of 60 ms, at a resolution of 120,000. Targeted MS2 scan spectra were acquired in the m/z range of 400-1000 using 16 m/z quadrupole isolation windows, AGC target of 1e6, at a resolution of 30,000, maximum injection time of 55 ms and loop count of 45. Higher-energy collision-induced dissociation fragmentation was performed in one-step collision energy of 27%. Electrospray voltage was static and capillary temperature was 275°C, with expected LC peak width of 30 sec. No sheath and auxiliary gas flow were used.

### Data-independent acquisition protein identification using DIA-NN

Raw files were searched in DIA-NN version 1.8 against a Uniprot *Mus musculus* database (containing 25,387 entries, downloaded 15/03/21) as well as a common contaminants database (Frankenfield et al., 2022), at 1% false discovery rate with default settings. *In silico* digest for spectral library generation used for library-free search.

### Proteomics data analysis

Proteins were first filtered to remove contaminants and those identified by only one unique peptide. For whole cell proteome analysis, Perseus V2.0.7.0 was used to generate heatmaps with hierarchical clustering and GO term enrichment according to the protocol outlined by Tyanova and Cox (Tyanova and Cox, 2018). Rstudio was used to log2 transform and median normalise the data, and determine which proteins showed significantly differential expression (adjusted p-value < 0.05 and fold change ≥ 2 or ≤-2) using the limma package with Benjamini-Hochberg correction used for multiple comparison testing (Ritchie et al., 2015). GSEA was performed using the clusterProfiler package with a minimum and maximum gene set size of 20 and 600, respectively. Benjamini-Hochberg correction was used for multiple comparisons with a p value cut off of 0.05. Both hallmark gene sets defined by the Molecular Signatures Database (MSigDB) and gene sets defined by the Kyoto Encyclopaedia of Genes and Genomes (KEGG) were used for Gene Set Enrichment Analysis (GSEA) analysis (Wu et al., 2021). STRING network analysis was performed with a minimum required interaction score of 0.4 using the following interaction sources: text mining, experiments, databases, and co-expression. Disconnected nodes were hidden from the visual network but included in the inbuilt enrichment analysis.

### Flow cytometry

Cells were plated in a V-bottom 96-well tissue culture plate (Cellstar) at 1×10^6^ cells per well in ice-cold Fluorescence-Activated Cell Sorting (FACS) buffer (1% BSA and 1% FBS in PBS, pH 7.2) and washed by centrifugation at 1,000 xg for 3 minutes. Cells were resuspended in 100 µL FACS buffer supplemented with Fc-γ receptor anti-CD16/CD32 antibodies (1:100, Thermo Fisher Scientific) and incubated on ice at 4°C for 30 minutes, to ensure Fc receptors were blocked. Cells were then washed and incubated with 100 µL FACS buffer containing the relevant surface protein targeted fluorophore-conjugated antibody (1:100, **Table 2.3**) on ice at 4°C for 30 minutes. Cells were fixed using 100 µL chilled 4% paraformaldehyde in PBS (Thermo Fisher Scientific) at room temperature for 20 minutes and subsequently washed with FACS buffer to remove excess paraformaldehyde. Samples were resuspended in 400 µl FACS buffer and analysed using a BD FACSymphony A5 flow cytometer (BD Biosciences). Gating and data analysis were performed either with unstained samples or fluorescence minus one control using FlowJo v10.8.1 (BD Biosciences).

**Table 0.**
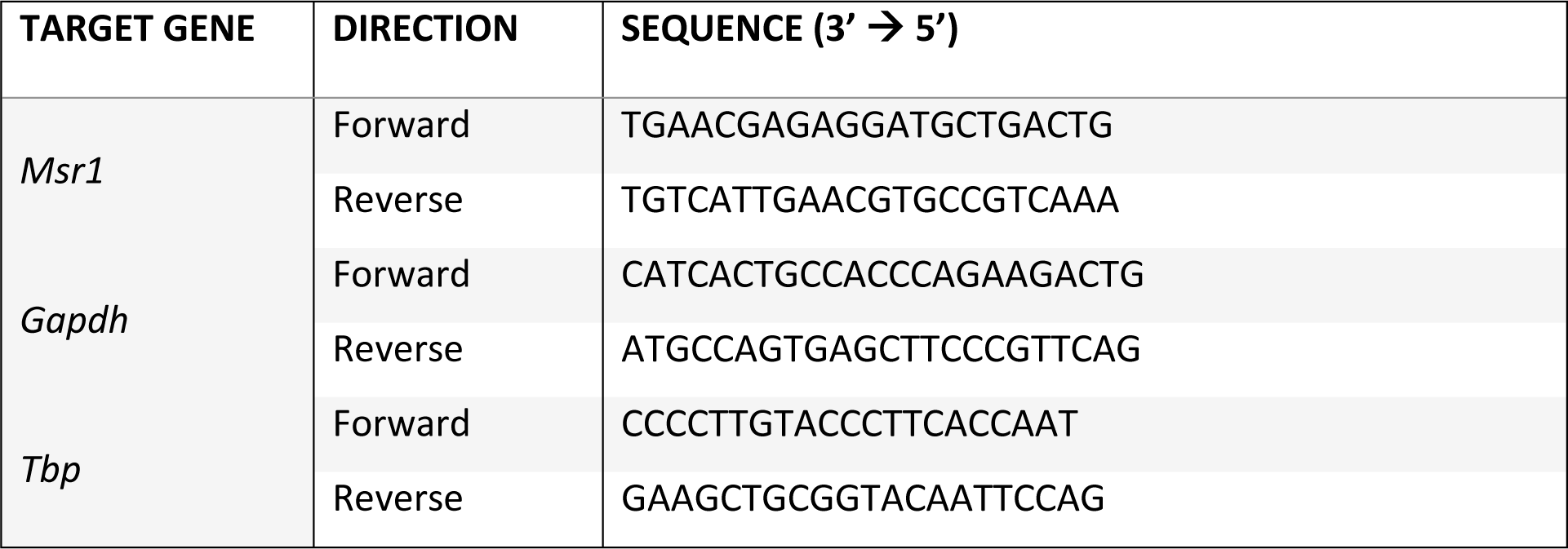
List of primers.

**Table 3.** Flow cytometry antibodies.

| ANTIBODY-CONJUGATE | SUPPLIER | CATALOGUE NO. | HOST SPECIES |
| --- | --- | --- | --- |
| F4/80- Alexa Fluor™ 488 | Invitrogen | 53-4801-82 | Rat |
| CD11B-BUV737 | Invitrogen | 367-0112-82 | Rat |
| CD11C-APC | Invitrogen | 17-0114-82 | Armenian hamster |
| MSR1-PE | Invitrogen | 12-2046-82 | Rat |

## Data Availability

The mass spectrometry proteomics data have been deposited to the ProteomeXchange Consortium via the PRIDE partner repository with the data set identifier: PXD051067. Reviewer account details: Username: Password: arFVjKcR

## Acknowledgements

We would like to thank Abeer Dannoura for technical support. This research was partly funded by a Wellcome Trust Investigator Award (215542/Z/19/Z) and an MRC DiMeN DTP Studentship to JG.

